# Structure of the *Macrobrachium rosenbergii* Nodavirus: A new genus within the *Nodaviridae?*

**DOI:** 10.1101/386888

**Authors:** Kok Lian Ho, Mads Gabrielsen, Poay Ling Beh, Chare Li Kueh, Qiu Xian Thong, James Streetley, Wen Siang Tan, David Bhella

**Affiliations:** Department of Pathology, Faculty of Medicine and Health Sciences, Universiti Putra Malaysia, 43400 UPM Serdang, Selangor, Malaysia.; CRUK Beatson Institute, Garscube Campus, Switchback Road, Glasgow, G61 1BD, Scotland UK.; Department of Microbiology, Faculty of Biotechnology and Biomolecular Sciences, Universiti Putra Malaysia, 43400 UPM Serdang, Selangor, Malaysia.; MRC - University of Glasgow Centre for Virus Research, Sir Michael Stoker Building, Garscube Campus, 464 Bearsden Road, Glasgow G61 1QH, Scotland, UK.; Institute of Bioscience, Universiti Putra Malaysia, 43400 UPM Serdang, Selangor, Malaysia

**Keywords:** Virus-like particles, insect cells, nodavirus, giant freshwater prawn, 3-dimensional structure, cryo-electron microscopy

## Abstract

*Macrobrachium rosenbergii* nodavirus (*Mr*NV) is a pathogen of freshwater prawns that poses a threat to food-security and causes significant economic losses in the aquaculture industries of many developing nations. A detailed understanding of the *Mr*NV virion structure will inform the development of strategies to control outbreaks. The *Mr*NV capsid has also been engineered to display heterologous antigens, thus knowledge of its atomic resolution structure will benefit efforts to develop tools based on this platform. Here we present an atomic-resolution model of the *Mr*NV capsid protein, calculated by cryogenic electron microscopy (cryoEM) of *Mr*NV virus-like particles (VLPs) produced in insect cells, and three-dimensional image reconstruction at 3.3 Å resolution. CryoEM of *Mr*NV virions purified from infected freshwater prawn post-larvae yielded a 6.6 Å resolution structure confirming the biological relevance of the VLP structure.

Our data revealed that unlike other known nodaviruses structures, which have been shown to assemble capsids having trimeric spikes, *Mr*NV assembles a T=3 capsid with dimeric spikes. We also found a number of surprising similarities between the *Mr*NV capsid structure and that of the *Tombusviridae*. 1. An extensive network of N-terminal arms lines the capsid interior forming long-range interactions to lace together asymmetric units. 2. The capsid shell is stabilised by three pairs of Ca^2+^ ions in each asymmetric unit. 3. The protruding spike domain exhibits a very similar fold to that seen in the spikes of the tombusviruses. These structural similarities raise questions concerning the correct taxonomic classification of *Mr*NV.

## Introduction

The giant freshwater prawn *Macrobrachium rosenbergii* is widely cultivated in tropical and subtropical areas for food. The global production of this species has increased dramatically from about 3,000 tons in 1980 to more than 220,000 tons in 2014 [1]. However, productivity is threatened by white tail disease (WTD), which is caused by *Macrobrachium rosenbergii* nodavirus (*Mr*NV). This often leads to 100% mortality rates in larvae and post-larvae of *Macrobrachium rosenbergii* [2]. The first *Mr*NV outbreak was reported in Pointe Noire, Guadeloupe in 1997, followed by China [3], India [4], Taiwan [5], Thailand [6], Malaysia [7], Australia [8] and recently in Indonesia [9]. To date, neither vaccine nor effective treatment is available to prevent or manage *Mr*NV outbreaks.

*Mr*NV has been classified within the *Nodaviridae* family of viruses. These non-enveloped viruses have bipartite, positive-sense, single-stranded RNA genomes that are packaged within T=3 icosahedral capsids. Presently this family has two established genera: *Alphanodavirus* and *Betanodavirus*. The former consists of insect-infecting nodaviruses such as Flock House virus (FHV), Pariacoto virus (PaV), black beetle virus (BBV), Nodamura virus (NoV), and Boolarra virus (BoV), while the latter contains fish-infecting nodaviruses such as Malabaricus grouper nervous necrosis virus (MGNNV), grouper nervous necrosis virus (GNNV) and striped jack nervous necrosis virus (SJNNV). Although *Mr*NV is classified within the *Nodaviridae*, amino acid sequence comparison revealed that its capsid protein shares low similarity (less than 20%) with other nodaviruses in the two established genera. Thus, it is ambiguous to group *Mr*NV in either one of these genera. Conversely, the amino acid sequence of *Mr*NV capsid protein shares approximately 80% similarity with that of *Penaeus vannamei* nodavirus (*Pv*NV). These two crustacean nodaviruses have therefore been proposed to be grouped into a new genus; *Gammanodavirus* [10].

Nodaviruses are very simple, their two, short genomic RNAs encoding three gene products. RNA 1 (3.2 kb) encodes the RNA-dependent RNA polymerase (RdRp) and the non-structural B2-like protein, while RNA 2 (1.2 kb) encodes the viral capsid protein (CP). The full-length *Mr*NV CP is a polypeptide of 371 amino acids. The N-terminal arginine rich region interacts with the RNA genome [11] while the C-terminal domain plays crucial roles in host cell attachment and internalization [12]. A nuclear localization signal (NLS) has also been identified at the N-terminus (amino acids 20-29) of CP. This has been shown to target the viral capsid to the nucleus of insect cells [13]. Further functional regions of CP have yet to be defined.

Several 3-dimensional (3D) structures of alpha- and beta-nodaviruses have been determined using both X-ray crystallography and cryo-electron microscopy [14-18]. These analyses have revealed several common features. Nodaviruses assemble T=3 icosahedral capsids; 180 CP protomers assemble such that the asymmetric unit comprises three identical capsid subunits in three quasi-equivalent positions termed A, B and C (here termed CP_A_, CP_B_ and CP_C_). To date, all alpha- and beta-nodaviruses have been found to have capsomeres that present a trimeric spike. The arginine-rich N-terminal region of the viral capsid protein interacts with the viral RNA segments [exemplified by PaV, PDB code: 1F8V [15]], leading to the formation of a dodecahedral RNA cage at the virion interior.

We have previously shown that recombinant CP of *Mr*NV produced using baculovirus expression in *Spodoptera frugiperda* (Sf9) cells, assembles into virus-like particles (VLPs) with a diameter of ˜40 nm [19]. We determined the intermediate resolution structure of the *Mr*NV capsid using cryo-electron microcopy and image reconstruction [20]. At this resolution, our reconstruction revealed distinct dimer-clustering of capsomeres in the T=3 *Mr*NV icosahedral capsid. Capsomeres were seen to form square, thin and blade-like spikes on the virion surface. All other nodaviruses have been shown to assemble with trimeric capsomers. Our structure therefore revealed a strikingly divergent morphology for *Mr*NV, lending weight to the proposed classification of *Mr*NV within a new genus of nodaviruses [20].

Here we present a high-resolution 3D reconstruction of the *Mr*NV VLP, solved at 3.3 Å resolution. From these data, we have constructed an atomic model of the *Mr*NV capsid protein. We show that the shell (S) domain of *Mr*NV CP possesses the canonical 8-stranded beta-barrel structure common to all nodaviruses. There are however striking structural similarities between the *Mr*NV capsid and those of members of the Tombusviridae. The protruding (P) domain, exhibits a similar fold to that which has been previously shown for tomato-bushy stunt virus (TBSV), although the spikes are narrower and are oriented quite differently in the two dimeric forms (AB and CC). CP-CP interactions in the S-domain are stabilised by coordinated Ca^2+^ ions. Protomers forming CC-dimers, located at the icosahedral two-fold symmetry axes of the capsid, possess an ordered N-terminal arm (NTA). This passes along the capsid interior forming inter-molecular interactions with neighbouring protomers to stabilise the capsid. Unlike the NTA of TBSV however, which folds back to form an additional β-strand in CP_C_ before going on to form a structure known as a β-annulus at the adjacent icosahedral three-fold axis, the NTA of the *Mr*NV CP_C_ crosses the icosahedral two-fold symmetry axis and inserts into a symmetry related CP_B_. The NTA then forms a β-annulus at the *next* nearest three-fold symmetry axis before continuing to pass along the capsid interior donating a strand to a β-sheet in a second neighbouring CP_B_ molecule.

We also present an intermediate-resolution structure of the authentic *Mr*NV virion, purified from infected larvae of *Macrobrachium rosenbergii*, revealing that the infectious virus exhibits an identical capsid structure to that which we have determined for the VLP. Our data present a detailed structural view of this economically important pathogen and raise questions concerning the taxonomic classification of both *Mr*NV and the related *Pv*NV.

## Materials and Methods

### Construction of pGEM-T TARNA2 plasmid encoding MrNV-CP

The gene encoding *Mr*NV-CP was amplified from plasmid pTrcHis2-TARNA2 [21]. The forward and reverse primers used to amplify the coding region were 5’-ATG GCC CTT AAC ATC A<u>CC ATG G</u>CT AGA GGT AAA CA-3’ (*Nco*I restriction site is underlined) and 5’-CTA TCG TCG GCA ATA ATT AAG G<u>CG AAT TC</u>G AAG CTT ACG T-3’ (*Eco*RI restriction site is underlined), respectively. PCR profile was denaturation at 95 °C for 3 minutes, followed by 35 cycles of i) denaturation at 95 °C for 30 s, ii) annealing at 59 °C for 30 s, iii) and extension at 72 °C for 1 minute. The final extension was performed at 72 °C for 10 minutes. The PCR products were excised and purified using the QIAquick Gel Extraction Kit (Qiagen, Germany). The purified DNA was ligated with the linearised pGEM-T vector (Promega, USA) and introduced into competent *Escherichia coli* DH5α cells. The transformants were plated on Luria Bertani (LB) agar plates containing ampicillin (100 μg/ml). Following an overnight incubation at 37 °C, single bacterial colonies were picked and cultured in LB broth. The orientation and nucleotide sequence of the DNA insert were confirmed by DNA sequencing.

### Mutagenesis of pFastBac-HTC plasmid

To produce the *Mr*NV-CP without a His-tag, the QuickChange II site-directed mutagenesis kit (Agilent Technologies, USA) was used to create an *Nco*I restriction nuclease cutting site in the pFastBac-HTC plasmid (Invitrogen, USA). The primers used for mutagenesis were 5’-CGG GCG CGG ATC TCG GTC CGA AA<u>C CAT GG</u>C GTA CTA CCA TCA CC-3’ and 5”-GGT GAT GGT AGT ACG <u>CCA TGG</u> TTT CGG ACC GAG ATC CGC GCC CG-3’, where the *Nco*I restriction cutting site is underlined.

### Construction of recombinant bacmid pFastBac^TM^ HT C-TARNA2

The pGEM-T TARNA2 plasmid and the mutated pFastBacHT-C plasmid were digested with *Eco*RI and *Nco*I, respectively. The digested products were purified using the QIAquick Gel Extraction Kit (Qiagen. Germany) and ligated together to produce the pFastBacHTC-TARNA2. This was introduced into competent *E. coli* DH10Bac cells (Invitrogen, USA) and plated on LB agar plates containing kanamycin (50 μg/ml), gentamicin (7 μg/ml), and tetracycline (10 μg/ml). White bacterial colonies containing the recombinant plasmid were selected and cultured in LB broth. The recombinant bacmid DNA was extracted and the presence of DNA insert was confirmed by PCR. The primers used in the PCR were pUC/M13 forward 5’-CCC AGT CAC GAC GTT GTA AAA CG-3’ and pUC/M13 reverse 5’-AGC GGA TAA CAA TTT CAC ACA GG- 3’.

### Preparation of recombinant baculovirus stock

Sf9 cells (8 × 10^5^ cells/well) in a 6-well plate were transfected with the recombinant bacmid pFastBacHTC-TARNA2 using Cellfectin II reagent. The transfected cells were incubated at 27°C for 72 hours. The cell culture medium was harvested by centrifugation at 500 × g for 5 minutes at 4 °C. The P1 baculovirus stock was amplified by infecting the Sf9 cells (2 × 10^6^ cells/mL) in serum free Sf-900 III SFM medium (Gibco®, USA). The infected cells were incubated at 27 °C for 72 hours. The P2 baculovirus stock was harvested by centrifugation at 500 × g for 5 minutes at 4 °C and stored at 4 °C for subsequent experiments.

### Expression and purification of recombinant MrNV capsid

Sf9 cells were cultured as suspension cells at 27 °C in a serum free Sf-900 III SFM medium to reach a cell density of 2 × 10^6^ cells/ml. Recombinant baculovirus stock [10% (v/v)] was added into the culture, which was further incubated for 4 days at 27 °C. The *Mr*NV capsid and the Sf9 cells were separated by centrifugation at 500 × g for 5 min at 4 °C. The *Mr*NV capsid was precipitated at 60% (w/v) ammonium sulphate saturation for 2 hours at 4 °C. The proteins were pelleted by centrifugation at 18, 000 × g for 20 min at 4 °C. The pellet was resuspended in HEPES buffer A (20 mM HEPES, 100 mM NaCl; pH 7.4) and dialysed in the same buffer overnight. The dialysed sample was purified by size exclusion chromatography (SEC) using a HiPrep™ 16/60 Sephacryl® S-500 HR column (GE Healthcare, USA), which was attached to a fast protein liquid chromatography (FPLC) system (Akta Purifier; GE Healthcare, USA). The purified protein was concentrated with a 100 kDa molecular cut-off centrifugal concentrator (Pall, USA) and the protein concentration was determined with the Bradford assay [22].

### Isolation of authentic MrNV virions from giant freshwater prawn larvae

Lysate of *Mr*NV-infected post-larvae was prepared according to published methods [23] with some modifications. Briefly, the infected post-larvae were homogenized in HEPES buffer B, (25mM HEPES, 150 mM NaCl; pH 7.4) and the homogenate was centrifuged at 6,000 × g for 10 minutes at 4 °C to remove large debris. The supernatant was further clarified by centrifugation at 12,100 × g for 30 minutes at 4 °C. The clarified supernatant was loaded onto a sucrose gradient [8-50% (w/v)] and centrifuged at 210,000 × g for 4.5 hours at 4 °C. Fractions (500 µl) were collected and analysed by SDS-PAGE and Western blotting. Fractions containing *Mr*NV were pooled and dialysed in HEPES buffer B. The purified *Mr*NV was concentrated by centrifugation using a centrifugal concentrator (molecular weight cut-off 10 kDa, Vivaspin Turbo 15, Sartorius, Germany). The final concentration of purified *Mr*NV was determined using the Bradford assay [22].

### Cryo-electron microscopy

Purified *Mr*NV VLPs (at approximately 0.2 mg/ml) or virions (at approximately 0.1 mg/ml) were prepared for cryogenic transmission electron microscopy using an FEI Vitrobot Mk IV. Particles were imaged on thin-continuous carbon films that had been applied to C-flat holey carbon support films (R1.2/1.3; Protochips). Four µl of VLP or virion preparation was loaded onto a grid for 1 minute, blotted for 4 seconds and plunged into liquid ethane. Vitrified samples were imaged at low-temperature (around 95 K) and under low electron dose conditions using a JEOL 2200 FS cryo-microscope operated at a nominal magnification of 50k× and an accelerating voltage of 200 kV. Frozen grids were held in a Gatan 626 cryo-stage. Images were recorded on a Direct Electron DE20 camera as 2 second movies at 20 frames per second and ~1.5 electrons/pixel/frame. The pixel size was 1.11 Å/pixel. To collect high-resolution data on *Mr*NV VLPs, grids were imaged at the electron bioimaging centre (eBIC),

Diamond Light Source (UK) using an FEI Titan Krios operated at 47,170× magnification. Images were recorded on a Gatan K2 BioQuantum energy-filtered direct detector camera operated in zero-loss imaging mode with a slit width of 20 eV. Five second exposures were recorded in electron counting mode, at a frame-rate of four frames per second and a dose rate of 1.8 electrons/pixel/frame. The pixel size was 1.06 Å/pixel.

### Three-dimensional image reconstruction

All image processing was performed using Relion 2.1 [24]. Image stacks of movie frames were motion-corrected using motioncor2 [25]. Defocus estimation was performed using GCTF [26]. For each data-set a small subset of particle images was manually picked and subjected to 2D classification to prepare a template for automated particle picking. Thereafter, particles were automatically picked for all motion-corrected micrographs. Individual particle images were extracted in 512^2^ pixel boxes. These data were processed to calculate 2D class averages, to select a data-set of the best particles and exclude erroneously picked images of frost, debris etc. 3D classification with imposition of icosahedral symmetry was used to further exclude data that did not yield high-quality reconstructions. In our study of *Mr*NV VLPs we used our previously calculated 3D reconstruction as a template for starting the classification. For authentic virions, we performed the first 3D classification analysis using a Gaussian sphere as the starting model to prevent model bias, as previously described [20]. Finalised data-sets were subjected to unsupervised 3D refinement. This was followed by masking and post-processing of reconstructions calculated from half-sets of data, to determine final resolutions and apply appropriate sharpening to the maps.

### Atomic model building

Atomic models were built from the high-resolution density maps using the CCP-EM suite of programmes [27] in particular COOT [28]. The model was refined using REFMAC [29] and PHENIX [30]. Validation was performed using MOLPROBITY [31]. Secondary structure assignment was performed using STRIDE (http://webclu.bio.wzw.tum.de/cgi-bin/stride/stridecgi.py) [32]. Density maps and atomic resolution models were visualised using UCSF Chimera [33]. Validation of metal ion assignment was performed using the ‘checkmymetal’ server (https://csgid.org/metal_sites) [34]. Contact interface analysis was performed using the PISA server (http://www.ebi.ac.uk/msd-srv/prot_int/cgi-bin/piserver) [35]. A 3D pairwise alignment of the *Mr*NV Cp P-domain structure and that of cucumber necrosis virus (PDB ID 4LLF) was performed using the FatCat server (http://fatcat.burnham.org) [36]. Protein topology diagrams were generated using Pro-Origami (http://munk.csse.unimelb.edu.au/pro-origami/porun.shtml) [37] and edited using Inkscape (https://inkscape.org/en/).

### Data Deposition

The cryoEM map of the *Mr*NV VLP was deposited in the electron microscopy databank with accession number EMD-0129. The cryoEM map of the *Mr*NV virion was deposited in the electron microscopy databank with accession number EMD-0130. The atomic coordinates for the asymmetric unit of the *Mr*NV VLP were deposited in the protein databank with accession number PDB-6H2B. The cryoEM image data for EMD-0129 are deposited in EMPIAR as motion-corrected single frame micrographs with accession number EMPIAR-10203.

## Results

### Cryo-EM of MrNV VLPs

To calculate an atomic model of the *Mr*NV capsid protein we sought to determine a high-resolution 3D reconstruction of the capsid. Frozen hydrated preparations of *Mr*NV VLPs were imaged in a FEI Titan Krios at the UK electron bioimaging centre (eBIC). A total of 2,459 cryomicrograph movies were recorded on a Gatan K2 BioQuantum direct detector (Fig. 1a). These data were processed to correct the effects of specimen movement and estimate the defocus in each micrograph. To calculate a template for automated particle picking, a dataset of approximately 1000 particles was manually picked and subjected to two-dimensional classification. Representative class-averages were selected and used to automatically pick particles for further analysis. A total of 60,939 putative particles were extracted from our motion-corrected micrographs and subjected to 2D classification. Class averages showing particle images with well-resolved structure were selected, reducing the data set to 56,762 particles. 3D classification was then used to select the best particles for inclusion in the final reconstruction, further reducing the dataset to 40,883 particles. Unsupervised refinement of the final dataset led to a reconstruction with an overall resolution of 3.3 Å (Fig. 1b-c and S1, Movie S1). The reconstructed density map closely matched our previously published 7 Å structure of the *Mr*NV VLP produced in *Sf*9 cells, showing pronounced blade-shaped dimeric spikes on the capsid exterior and a dodecahedral cage of RNA density within the particle. A cross-section through the reconstructed density revealed that the S-domains of the VLP were sharply resolved, while the P-domains were less well-defined, having weaker fuzzier density (Fig. 1b). Local resolution analysis confirmed this, revealing that much of the S-domain was solved to 3.2 Å resolution, while the tips of the dimeric capsomeres were poorer than 4 Å resolution (Fig. 1d). Local resolution filtering and sharpening was applied with a B-factor of - 140 Å^2^ to generate a density map that was suited for high-resolution model building.

**Figure 1.**
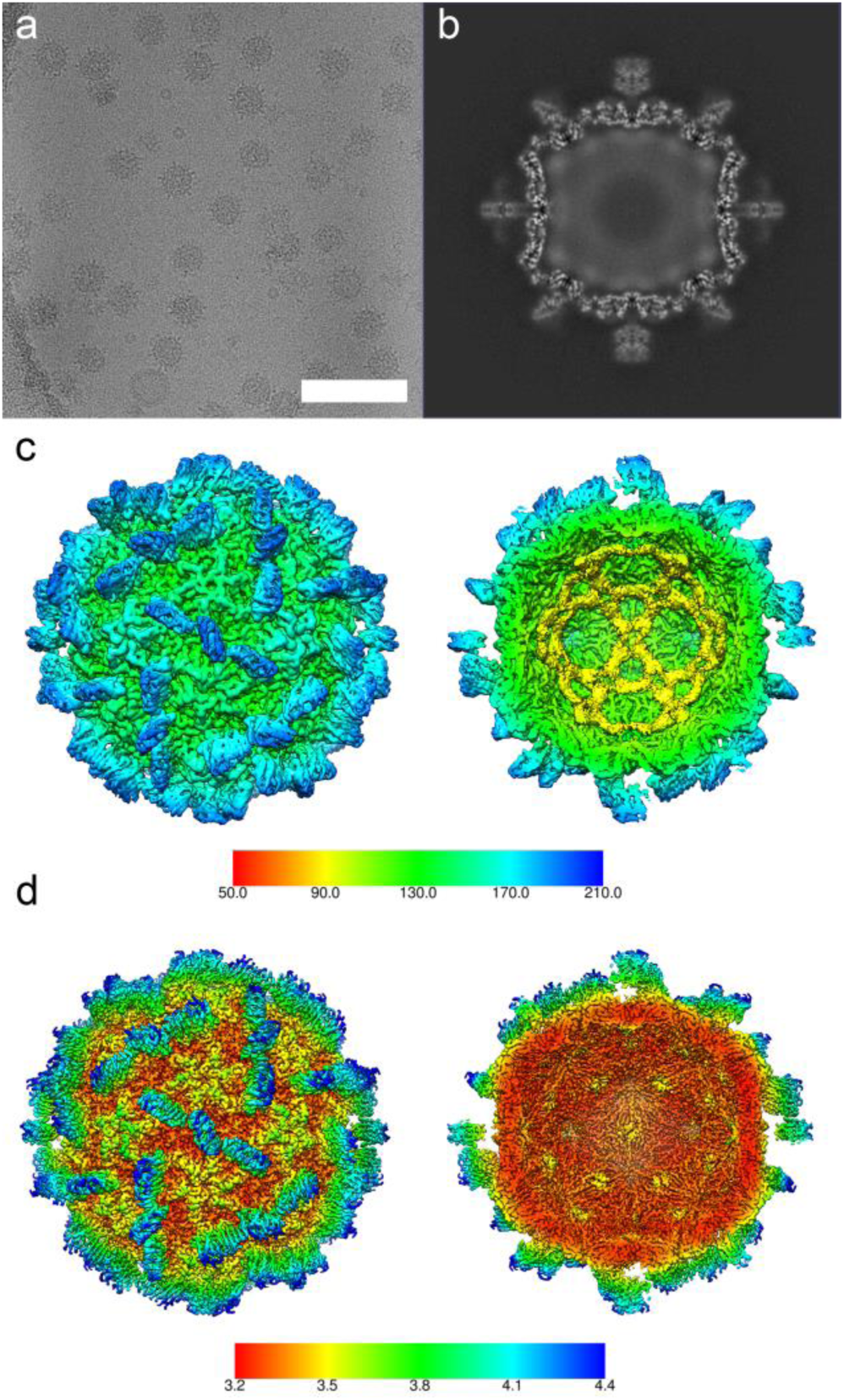
CryoEM and 3D reconstruction of MrNV VLPs. CryoEM of *Mr*NV VLPs revealed particles that were 40nm in diameter with pronounced spikes on their outer surface (a, scale bar = 100nm). A central section through the icosahedral reconstruction of the *Mr*NV VLP reveals a sharply defined capsid shell containing fuzzy density that we attribute to packaged nucleic acids. The P-domain spikes were less well resolved in comparison to the shell, most likely a consequence of flexibility (b). An isosurface view of the reconstruction is shown coloured by radius. A cutaway view reveals that the internal density forms a dodecahedral cage, consistent with that described for other nodaviruses (c). The sharpened map is also presented, coloured according to resolution, as both external and cutaway views (d). Sharpening of cryoEM maps down-weights lower resolution information to reveal the fine structural details of the map, such as amino acid sidechains. Poorly defined features such as the packaged nucleic acids are often lost upon sharpening, as no high-resolution information is present.

### An atomic model of the MrNV VLP asymmetric unit

The asymmetric unit of the T=3 *Mr*NV capsid comprises three copies of *Mr*NV-CP; CP_A_, CP_B_ and CP_C_. We have previously shown that the P domains assemble to form dimeric spikes; AB dimers arranged about the five-fold symmetry axes and the CC dimers located at the two-fold symmetry axes. We set out to build the sequence of the *Mr*NV capsid protein (Fig. S2) into our density map to produce an atomic model of the *Mr*NV capsid protein for each quasi-equivalent position. As a starting point, we docked a homology model into our density map [12, 20]. Overall this model fitted poorly within the reconstructed density map, with the exception of two regions between amino acid residues 104-135 and 232-243. A model for the S-domain of CP_A_ was therefore manually built and refined from this starting point. This partial model was then docked to CP_B_ and CP_C_ and further edited and refined, leading to reliable models for the S-domains at each quasi-equivalent position. Interestingly, our density map presented density consistent with the presence of two metal ions per capsid protein. Based on the surrounding residues and distances to coordinating atoms, we have modelled these as calcium ions (discussed below). In our 3D reconstruction, density for the S-domain was very well resolved. This allowed us to model this region to a high degree of confidence and with relative ease.

The P-domains were however rather less well defined and presented a more challenging task, particularly at the distal tips of the dimeric-spikes. CP_B_ was found to be the best resolved as judged by continuity of density, while CP_C_ appeared the least well resolved. Throughout the amino acid sequence of the P-domain there are bulky amino acid side-chains that gave confidence in our interpretation of the map. Following several rounds of manual editing and refinement a model was achieved for the full asymmetric unit that matched our density map and had reasonable geometry (Fig. 2, Movie S2, Table S1).

**Figure 2.**
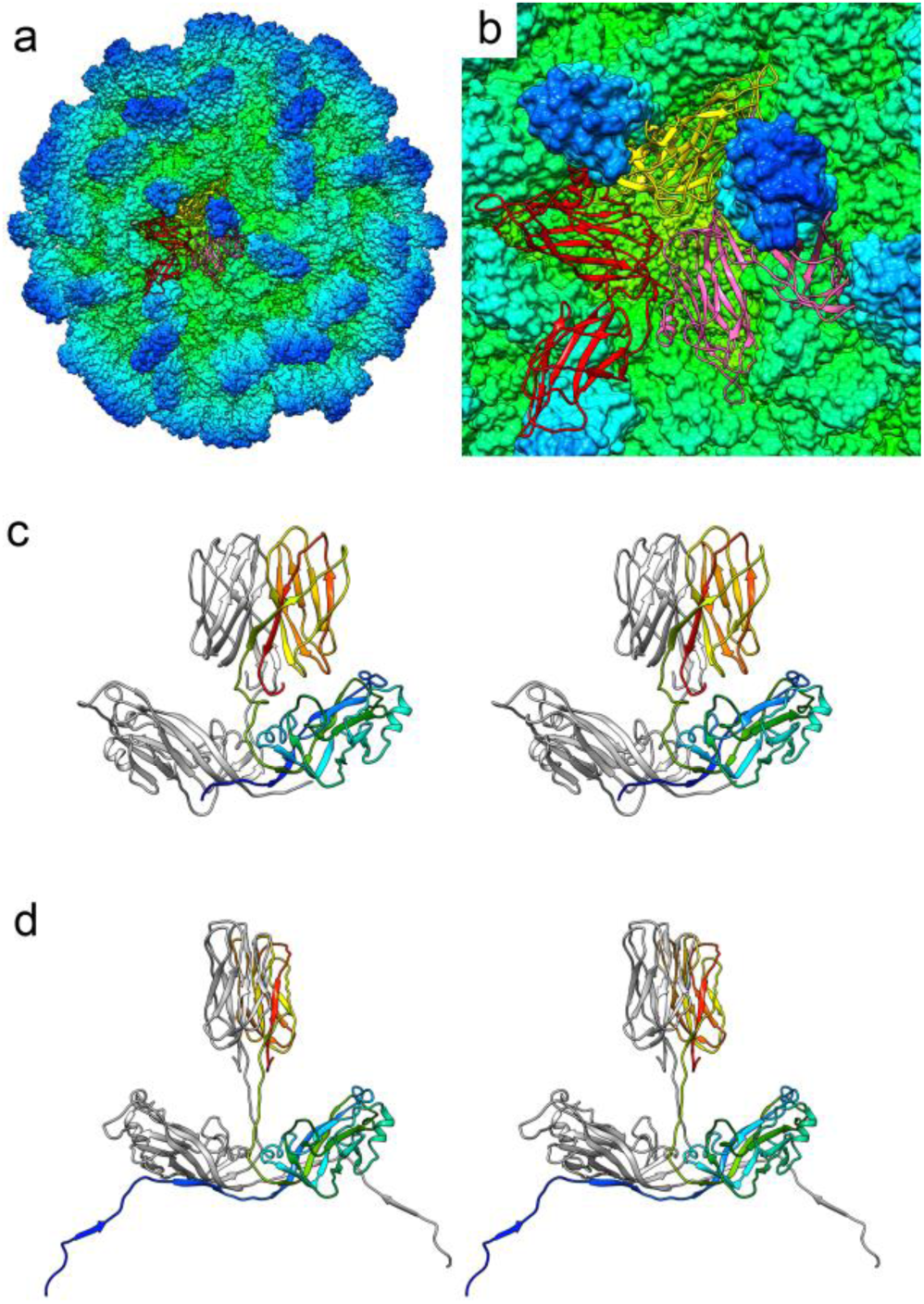
Atomic model of the MrNV capsid. A solvent excluded surface of the entire capsid is shown, coloured by radius (see Fig. 1c for key). A single asymmetric unit comprising three copies of CP – CP_A_ (red), CP_B_ (yellow) and CP_C_ (pink) is shown as a ribbon diagram (a), close up view (b). Wall-eyed stereo paired views of each dimer are shown (AB-dimer (c), CC-dimer (d)). In (c) CP_B_ is presented in rainbow colour scheme, while CP_A_ is coloured grey. In (d) one CP_C_ chain is shown in rainbow colour scheme, while the other is grey.

### A β-annulus motif formed of multiple N-terminal arms of chain C

The N-terminal regions of CP that include the RNA binding sites (amino acid residues 21-29) were not resolved for any of the chains (CP_A_, CP_B_ or CP_C_) in our density map. For CP_A_, we have successfully modelled amino acid residues 56-371, while for CP_B_ we were able to build amino acid residues 55-371. CP_C_ has a well-resolved NTA that allowed modelling from amino acid 31. Interestingly the CP_C_ NTA was found to form extensive contacts with symmetry related CP_C_ molecules, forming a network that crosses the capsid interior and is reminiscent of the NTAs previously described for several tombusviruses (Fig. 3, Movie S3). The CC dimer interface lies at the icosahedral two-fold axis. Each CP_C_-NTA emerges from the S-domain close to this symmetry axis and interacts with two CP_B_ protomers, donating β-strands to β-sheets within the CP_B_ S-domains. The CP_C_-NTA extends across the CC-dimer (and icosahedral) two-fold symmetry axis and inserts into the first CP_B_ subunit which lies adjacent to the symmetry related CP_C_ subunit (Fig. 3c-d). Moving from the C-terminus to the N-terminus, the NTA then crosses the adjacent icosahedral three-fold axis and inserts into the next nearest CP_B_ subunit, donating a second β-strand to the β-sheet comprising that molecule and a symmetry related CP_C_-NTA (Fig. 3e-f). This inter-digitated arrangement of NTA’s extending from capsid proteins at the C-position was first described for TBSV [38] and termed a β-annulus owing to the manner in which CP_C_-NTAs wrap around each other at the icosahedral three-fold axes. Nevertheless, in the case of *Mr*NV the lacing together of CP_C_ molecules is more extensive, as the NTA does not fold back on the CP_C_ to emerge from the S-domain at the nearest three-fold axis, and there form the β-annulus structure (as it does in TBSV). Rather it crosses the *two*-fold axis of the CC-dimer and then inserts into two CP_B_ molecules arranged about the *opposite* three-fold axis, where the β-annulus is formed. The last resolved N-terminal residue (Pro31) lays under the next neighbouring two-fold axis. This is related to the originating two-fold by a counter-clockwise rotation of 120^°^ about the three-fold axis of the β-annulus (viewed from the capsid exterior). Although it is not resolved, the arginine-rich putative RNA binding site (amino acids 21-29), must therefore be located proximal to the two-fold symmetry axes for the CP_C_-chains.

**Figure 3.**
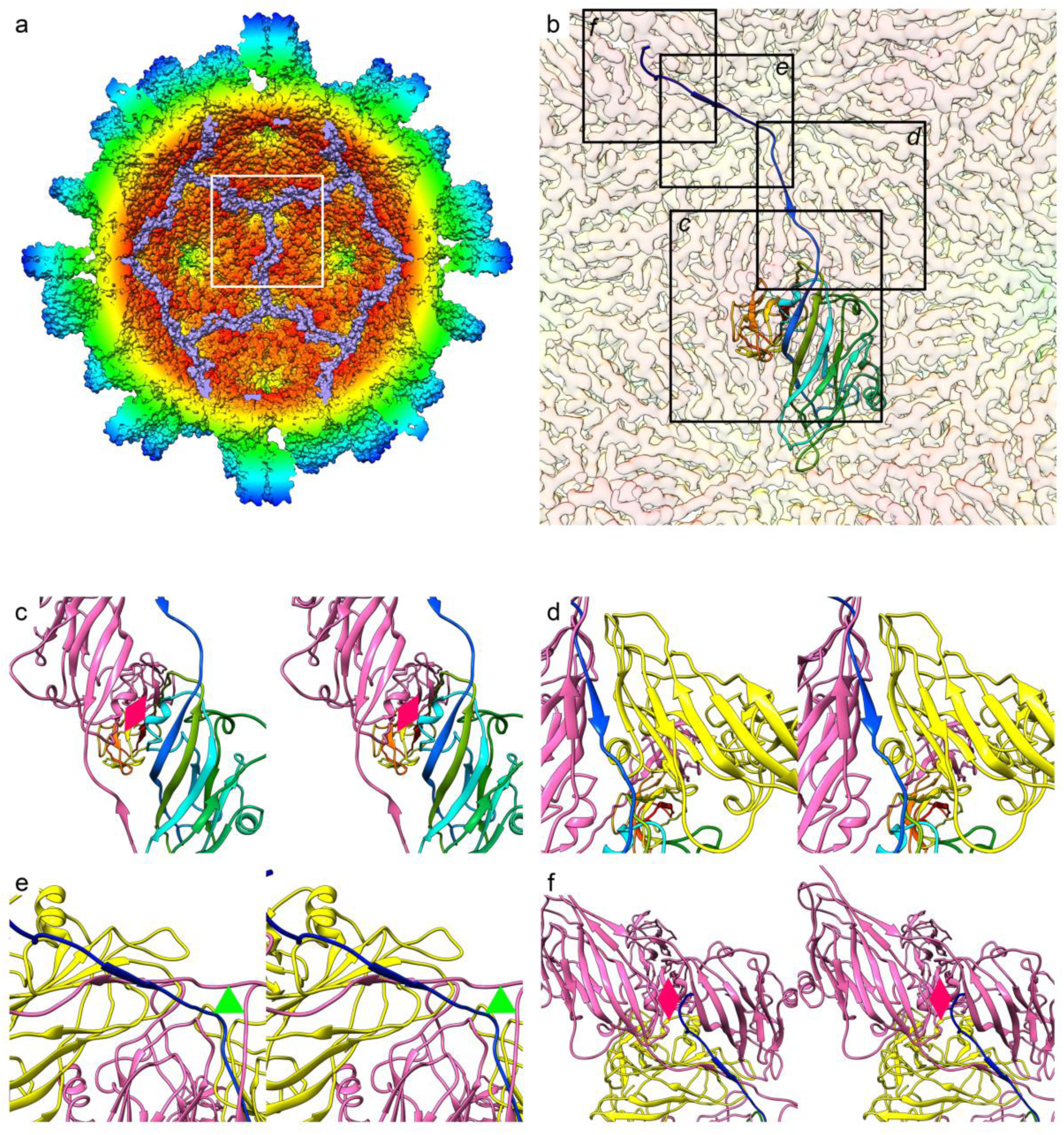
The CP_C_ NTA forms an extensive network at the capsid interior that includes α β-annulus motif. A solvent excluded surface view of the *Mr*NV capsid is shown, coloured by radius and clipped to reveal the capsid interior. The CP_C_ NTAs are coloured mauve to highlight the extended network formed (a). The white frame highlights the view presented in (b), in which the cryoEM map is shown as a transparent isosurface, with a single CP_C_ molecule presented as a ribbon diagram with rainbow colouring. Black frames highlight the views presented in subsequent figures (Fig 3c-f). (c) A wall-eyed stereo pair view of a CC-dimer, from the capsid interior shows the NTA emerging from the S-domain β-jelly roll to cross the icosahedral two-fold symmetry axis (pink diamond). (d) Moving from C-to N-termini, the NTA runs between the symmetry related CP_C_ (pink) and an adjacent CP_B_ (yellow) donating a strand to a β-sheet within the CP_B_. This sheet also includes a β-strand donated by a symmetry related CP_C_-NTA (pink). (e) The NTA then forms a β-annulus at the icosahedral three-fold axis (green triangle) and donates a strand to a β-sheet within a second CP_B_ (yellow), which similarly also includes a β-sheet from a symmetry related CP_C_ (pink). Thus CP_C_-NTAs are interdigitated between other CP_C_-NTAs and CP_B_ S-domains. (f) The N-terminal 29 residues of CP_C_ are not resolved in our cryoEM map, however the last visible CP_C_-NTA density terminates beneath a symmetry-related icosahedral two-fold symmetry axis (pink diamond), suggesting that the RNA binding site is proximal to this region.

### Coordinated metal ions stabilise CP-CP interactions in the S-domain

The *Mr*NV-CP S-domain comprises residues 62-242 and forms the contiguous shell of the capsid. The T=3 assembly is made up of 180 copies of the canonical 8-stranded anti-parallel β-barrel fold, known as the β-jelly roll. This is commonly seen in positive sense RNA containing viruses including both the nodaviruses and tombusviruses. Another interesting parallel between the structure of *Mr*NV and the tombusviruses is the presence of coordinated metal ions at the interface between CP subunits (Fig. 4, Movie S4). X-ray crystallographic difference mapping of TBSV following EDTA treatment identified two divalent cation binding sites at the interface between capsid proteins within the asymmetric unit, that were modelled as Ca^2+^ [39]. Chelation followed by a rise in pH (>7.0) has been shown to cause a structural transition to a ‘swollen’ state in these virions, indicating that these divalent cations play a role in virion stabilisation or possibly control of uncoating. Based on the striking similarity in the locations of these putative metal ions in our data compared to those previously published for TBSV (PDB ID 2TBV), and the surrounding residues, we have modelled these putative metal ions as calcium (Fig. 4b).

**Figure 4.**
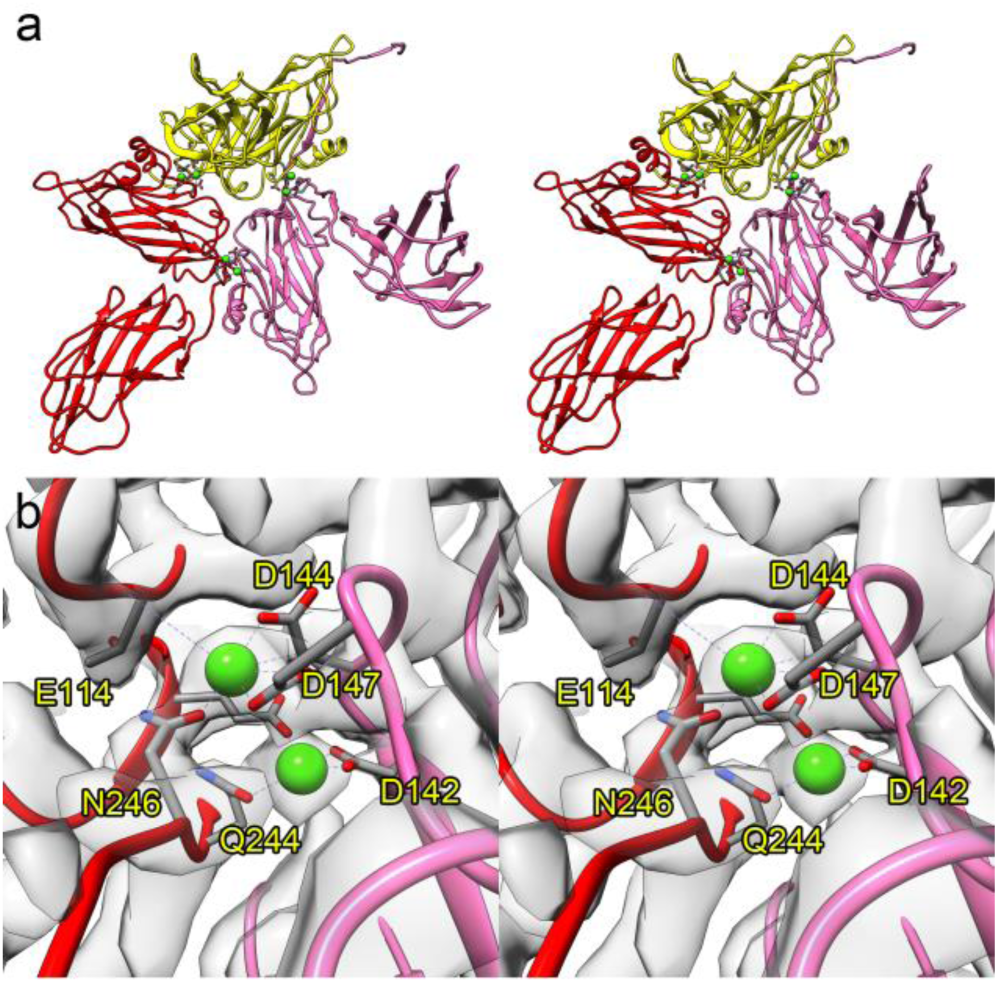
Metal ions stabilise the MrNV asymmetric unit. Within our cryoEM map we saw density consistent with the presence of coordinated metal ions. A wall-eyed stereo paired view of a ribbon diagram of the asymmetric unit is shown with chains coloured A-red, B-yellow and C-pink. Green spheres highlight the positions of the putative metal ions, which we have modelled as Ca^2+^ (a). A close-up view of a pair of metal ions at the interface between chains A and C is also shown in stereo (b). The amino acid residues that make up the coordination spheres are labelled. The cryoEM density map is also shown as a transparent isosurface.

### A flexible hinge leads to differently oriented P-domain dimer spikes

We have previously noted the striking differences in the orientations of the P-domain dimer spikes, relative to the underlying capsid shell, between AB and CC dimers. The CC-dimer spike is rotated approximately 85° counter-clockwise relative to that of the AB-dimer (viewed from the capsid exterior). Moreover, CC-dimer P-domains are raised from the capsid surface upon legs of density, whereas the AB P-domains sit closer to the capsid shell and are tilted towards their nearest two-fold symmetry axis. Our atomic model of the *Mr*NV VLP reveals the reason for the substantial differences in pose of these two capsomere forms. There is a large linker region between the S and P domains between amino acid residues 241 and 258. In the CC-dimer, this linker emerges from the S-domain β-jelly roll and forms a straight leg that is normal to the capsid surface (Fig. 2e). The inter-domain linker in CP_A_ and CP_B_ on the other hand has two bends, one between residues Pro247 and Pro249, which causes the linker to make a right-angled turn, and another at Ile252-Gln254 which likewise causes a right-angled turn, restoring the path of the linker to its original radial orientation (Fig. 2c). The twist in the linkers at the AB-dimer, induced by these turns, therefore accounts for the major differences in the orientations of the two types of spike. Interestingly, although we previously noted that the CP_B_ P-domain was more closely apposed to the S-domain than CP_A_, our model does not show any contacts. The AB P-domain’s orientation is defined by interactions with the AB linker region and the CP_C_ P-domain (Movie S5).

*Inter-capsomere contacts in the P-domain lead to the formation of a blade-like superstructure* CC-dimer spikes are less well-resolved in our cryoEM map than those of the AB-dimers. This is to be expected given the manner in which the AB-dimer spike is stabilised through interactions in the AB linker. In contrast, the P-domains of CP_C_ stand on extended polypeptide legs that may not offer the same support. The CC spike is instead stabilised by contacts between the P-domains of CP_B_ and CP_C_. AB spikes act as buttresses to the CC capsomere through polar interactions between amino acid residues 270 to 276 of CP_B_ and 307 to 317 of CP_C_ (Fig. 5, Movie S5). This gives rise to the formation of a blade-like superstructure that lays across the two-fold symmetry axis and comprises one CC-dimer and two AB-dimers.

**Figure 5.**
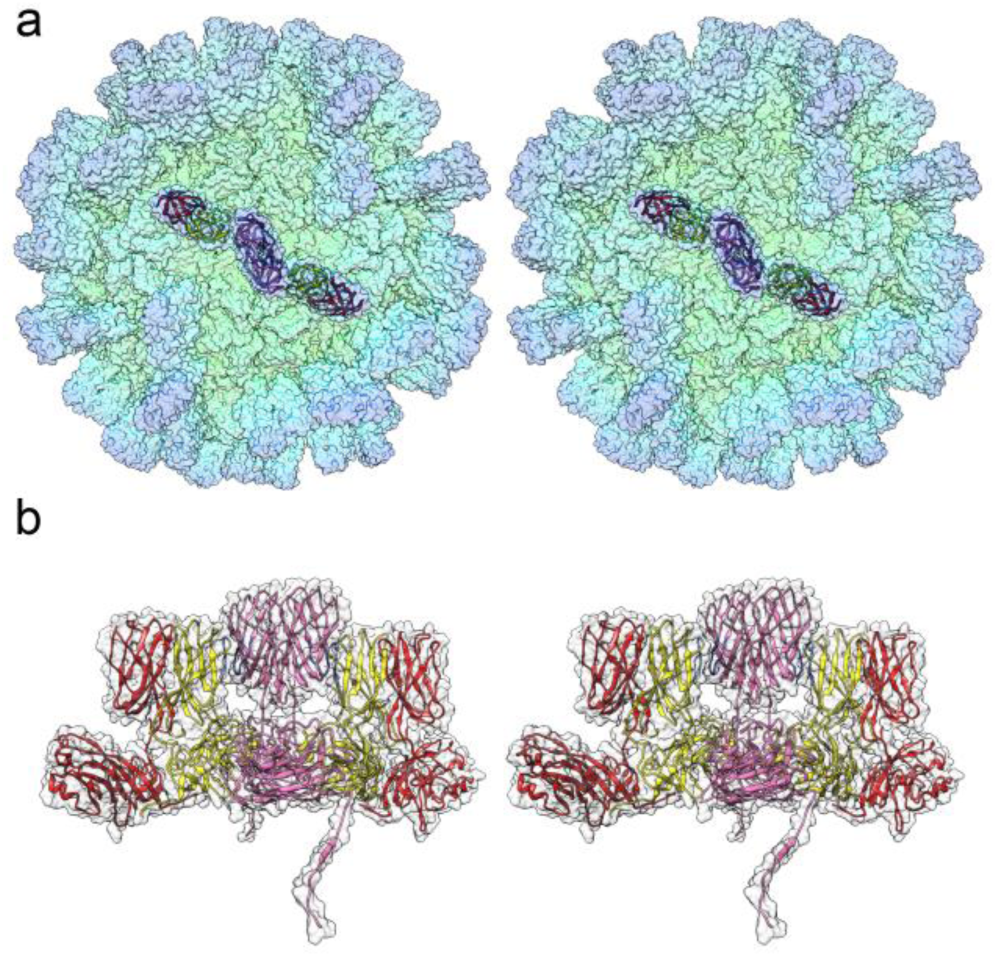
A superstructure comprising the P-domains of two AB-dimers and one CC-dimer. A transparent solvent excluded surface of the *Mr*NV capsid is presented as a stereo-paired view (a). Six P-domains are shown as ribbon diagrams to highlight the formation of a superstructure that lays across the icosahedral two-fold symmetry axis. A side-view of the superstructure shows how P-domain AB-dimers act as buttresses to stabilise the CC-dimer spike. The contact residues in CP_B_ (yellow) and CP_C_ (pink) are highlighted in shades of blue.

### The MrNV CP P-domain fold closely resembles that of the tombusviruses

It is noteworthy that the previously described homology model for the *Mr*NV CP structure [12] was based on the capsid protein of a tombusvirus: cucumber necrosis virus (CNV - PDB 4LLF [40]), rather than other known nodavirus structures. Although the homology model was a poor match for our cryoEM density map, our analysis has confirmed the hypothesised dimer-clustered T=3 icosahedral capsid structure. We have also discovered that like tombusviruses, *Mr*NV CP proteins bind metal ions to stabilise the capsid asymmetric unit. Close inspection of the fold of the *Mr*NV P-domain also reveals an unexpected structural homology between this nodavirus and the tombusviruses (Fig. 6). Tombusvirus P-domains have been shown to comprise a 10-stranded antiparallel β-barrel made up of two β-sheets annotated as BAJEHG and CDIF (Fig. 6a,d). Secondary structure assignment in the P-domains of the *Mr*NV structure was challenging, owing to the poorer resolution in this region. Nonetheless we have identified a similar β-barrel motif composed of nine strands arranged into two β-sheets annotated as AJEH2 and DIFGH1, based on a 3D pairwise alignment of the P-domains for CP_B_ of CNV and *Mr*NV (Fig. 6b,c, S3).

*Authentic MrNV virions assemble capsids indistinguishable from VLPs* To ensure that our atomic resolution model of the *Mr*NV capsid is an accurate description of the authentic virion, we determined the structure of purified virions at intermediate resolution. Virions were purified from homogenised, *Mr*NV infected post-larvae and imaged in a JEOL 2200 FS cryomicroscope. 7236 particle images were extracted from 263 micrographs and subjected to 2D and 3D classification. This led to the definition of a final dataset of 3931 particles that were further refined to produce the final reconstruction at 6.6 Å resolution (Fig. 7). At this resolution, the map appears identical to that of the *Mr*NV VLP in all respects (compare Fig. 7a and 1c). Furthermore, the packaged RNA shows very similar, albeit noisier, structure to the previously described dodecahedral cage. Thus, we conclude that our high-resolution model is an accurate representation of the structure of the authentic *Mr*NV virion.

**Figure 6.**
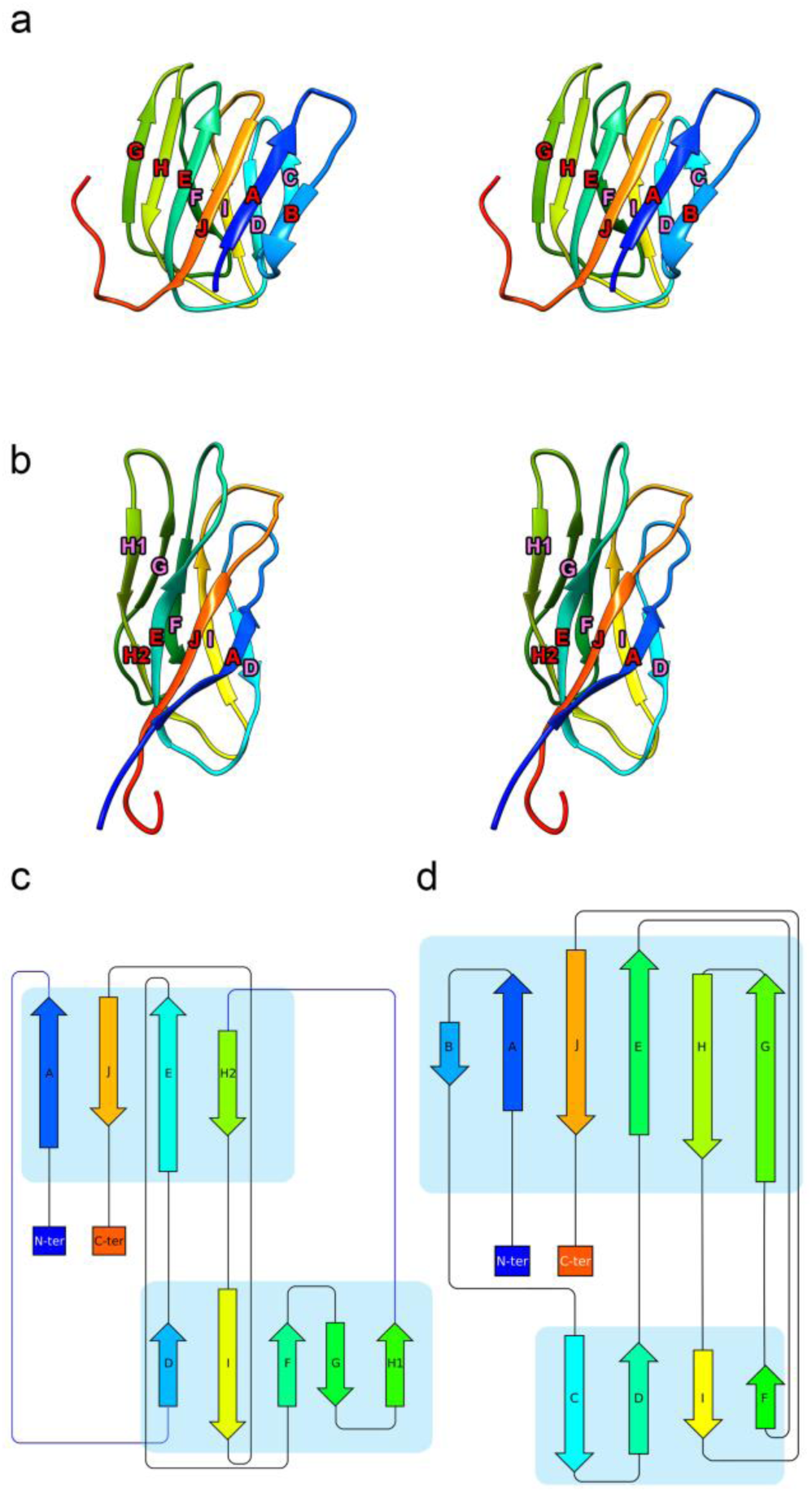
The MrNV P-domain has a similar fold to that of the tombusvirus cucumber necrosis virus. A stereo-paired view of the cucumber necrosis virus CP_B_ P-domain (PDB 4LLF) is presented as a ribbon diagram with rainbow colouring (a). The diagram is annotated to indicate successive β-strands from the N-to C-termini that together make up a ten-stranded antiparallel β-barrel. A similar motif is seen in the *Mr*NV P-domain, which is likewise presented and annotated (b). Protein topology diagrams of CNV CP_B_ P-domain (c) and *Mr*NV CP_B_ P-domain (d) present a simplified view to highlight the similarities of the P-domain folds in these two viruses.

**Figure 7.**
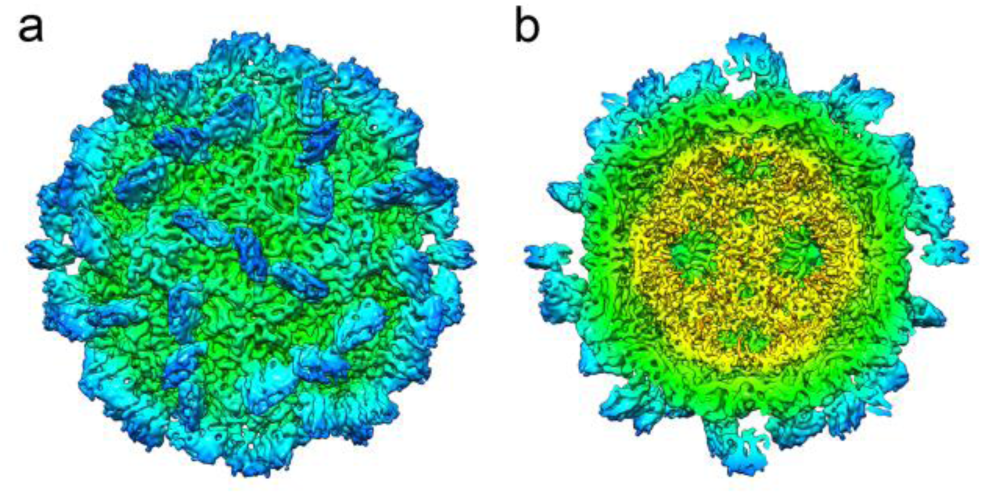
The authentic MrNV virion has an identical capsid structure to that of the VLP. CryoEM was used to calculate an intermediate-resolution 3D reconstruction of the authentic *Mr*NV virion, purified from infected *Macrobrachium rosenbergii* post-larvae. This revealed a structure that was indistinguishable from the VLP reconstruction (a), including the internal dodecahedral cage density, that we attribute to viral genome (b).

## Discussion

We have previously described the intermediate resolution structure of VLPs generated following recombinant expression of the *Mr*NV CP. This revealed a surprising divergence from known nodavirus structures [20]. The *Mr*NV T=3 icosahedral capsid was seen to assemble with dimeric rather than the usual trimeric capsomeres. Consistent with previously published nodavirus structures, we found that the *Mr*NV VLP exhibited density suggestive of packaging of nucleic acids – most likely the cognate mRNA. We also observed a surprising difference in the orientations of the P-domain spikes between the two classes of dimer (AB and CC).

Here we have extended this study, using cryoEM to calculate a 3D reconstruction of the *Mr*NV VLP at near-atomic resolution. This has allowed us to build an atomic-model of the capsid’s asymmetric unit. Our model reveals that *Mr*NV, like most small icosahedral positive-sense RNA viruses adopts the β-jelly roll fold in the S-domain. We identified major differences between AB- and CC-dimers, in the linker region that connects the S and P domains, accounting for the radically different orientations of their respective P-domains.

### Taxonomy of MrNV

Beyond providing a detailed description of the structure of *Mr*NV, our data revealed startling similarities between the *Mr*NV capsid structure and those of tombusviruses. We found that the *Mr*NV capsid’s asymmetric unit is stabilised by 6 Ca^2+^ ions, in a manner highly reminiscent of that seen in tomato bushy stunt virus. Furthermore, the CP_C_ NTA was found to form an extensive network at the capsid interior that involved an inter-digitated structure known as a β-annulus. This motif is also a feature of the tombusviruses. Finally, we found that the fold of the P-domain consisted of a β-barrel that also bore a close resemblance to the P-domain structure of the tombusviruses.

The divergence of amino acid sequence and distinct structural features of *Mr*NV compared with other nodaviruses led us to propose that *Mr*NV along with the related *Pv*NV might be classified into a new genus *Gammanodavirus* [10, 20]. Our more detailed description of the structure of the *Mr*NV capsid now adds to the weight of evidence arguing for close examination of the correct taxonomic classification of these viruses. Nodaviruses are characterised as small positive-sense RNA containing viruses, having bipartite genomes, that infect fish and invertebrates. Tombusviruses are plant-viruses that are classified on the nature of their RNA polymerases but are also seen to have consistent capsid structures. Thus, *Mr*NV having features of both virus families poses a conundrum with respect to its taxonomic status.

### P-domain structure and function

It has been demonstrated that the C-terminal domain of *Mr*NV is important for virus attachment and entry. In particular deletion of the last 26 amino acid residues substantially reduced infectivity. Inspection of the P-domain structure however, strongly suggests that deletion of this region is liable to significantly disrupt the β-barrel, as it would remove two strands from the centre of each β-sheet (Fig. S4). Thus, while it seems likely that the receptor binding site is within the P-domain, it may not be limited to the last 26 amino acid residues (345-371).

We have previously demonstrated insertion of heterologous epitopes into the *Mr*NV capsid structure at the C-terminus, such as the ‘a’ determinant of hepatitis B virus surface antigen (HBsAg) [41] and the ectodomain of matrix 2 protein (M2e) of influenza A virus [42]. Both epitopes were shown to be displayed on the surface of VLPs. Thus, *Mr*NV VLPs present an attractive platform for antigen display. Our structure of *Mr*NV CP now allows us to refine the placement of foreign epitopes. Indeed, each of the four loops on the outer surface of the P-domain (amino acid residues 268-275, 296-303, 322-326 and 350-355) represent potentially improved targets for further insertions. Combined with the capacity to package nucleic acids *Mr*NV VLPS may therefore prove to be a useful tool for both vaccine and DNA/RNA delivery.

### An economically important pathogen of freshwater prawns

*Mr*NV threatens livelihoods and food security in developing nations. Our atomic-resolution model of the *Mr*NV capsid provides insights into the fundamental biology of this important pathogen, highlighting features that may prove important in our understanding of virus assembly or entry, such as the presence of metal-ions that stabilise the asymmetric unit, and the structure of the receptor-binding P-domain. Such detailed understanding of the capsid structure provides a platform for the development of interventions to control or prevent disease outbreaks in the future.

## Acknowledgements

The authors wish to thank Diamond Light Source for access to the Cryo-EM facilities at the UK national electron bio-imaging centre (eBIC), proposal EM16810, funded by the Wellcome Trust, MRC and BBSRC. We thank Daniel Clare for his expert assistance during data collection. KLH was supported by Research Management Centre, Universiti Putra Malaysia (UPM). PLB and CLK were supported by the Graduate Research Fellowship, UPM and MyBrain scholarship, the Ministry of Higher Education, Malaysia. This work was funded by UPM Putra Grant (Project number: GP-IPS/2016/9509400) and the United Kingdom Medical Research Council (MC_UU_12014/7).

